# Novel drivers of virulence in *Clostridioides difficile* identified via context-specific metabolic network analysis

**DOI:** 10.1101/2020.11.09.373480

**Authors:** Matthew L Jenior, Jhansi L Leslie, Deborah A Powers, Elizabeth M Garrett, Kimberly A Walker, Mary E Dickenson, William A Petri, Rita Tamayo, Jason A Papin

## Abstract

The pathogen *Clostridioides difficile* causes toxin-mediated diarrhea and is the leading cause of hospital-acquired infection in the US. Due to growing antibiotic resistance and recurrent infection, targeting *C. difficile* metabolism presents a new approach to combat this infection. Genome-scale metabolic network reconstructions (GENREs) have been used to identify therapeutic targets and uncover properties that determine cellular behaviors. Thus, we constructed *C. difficile* GENREs for a hyper-virulent isolate (str. R20291) and a historic strain (str. 630), validating both with *in vitro* and *in vivo* datasets. Growth simulations revealed significant correlations with measured carbon source usage (PPV ≥ 92.7%), and single-gene deletion analysis showed >89.0% accuracy. Next, we utilized each GENRE to identify metabolic drivers of both sporulation and biofilm formation. Through contextualization of each model using transcriptomes generated from *in vitro* and infection conditions, we discovered reliance on the Pentose Phosphate Pathway as well as increased usage of cytidine and N-acetylneuraminate when virulence expression is reduced, which was subsequently supported experimentally. Our results highlight the ability of GENREs to identify novel metabolite signals in bacterial pathogenesis.

**Importance:** *Clostridioides difficile* is a Gram-positive, sporulating anaerobe that has become the leading cause of hospital-acquired infections. Numerous studies have demonstrated the importance of specific metabolic pathways in aspects of *C. difficile* pathophysiology, from initial colonization to regulation of virulence factors. In the past, genome-scale metabolic network reconstruction (GENRE) analysis of bacteria has enabled systematic investigation of the genetic and metabolic properties that contribute to downstream virulence phenotypes. With this in mind, we generated and extensively curated *C. difficile* GENREs for both a well-studied laboratory strain (str. 630) as well as a more recently characterized hyper-virulent isolate (str. R20291). *In silico* validation of both GENREs revealed high degrees of agreement with experimental gene essentiality and carbon source utilization datasets. Subsequent exploration of context-specific metabolism during both *in vitro* growth and infection revealed consistent patterns of metabolism which corresponded with experimentally measured increases in virulence factor expression. Our results support that differential *C. difficile* virulence is associated with distinct metabolic programs related use of carbon sources and provides a platform for identification of novel therapeutic targets.

## Introduction

*Clostridioides difficile* is a Gram-positive, sporulating anaerobic bacterium that remains a critical problem in healthcare facilities across the developed world (1, 2). Susceptibility to *C. difficile* infection (CDI) is most frequently preceded by exposure to antibiotic therapy (3). While these drugs are life-saving they also diminish the abundance of other bacteria in the microbiota, altering the metabolic environment of the gut, leaving it susceptible to colonization by *C. difficile* (4-6). Recently, we observed that *C. difficile* adapts transcription of distinct catabolic pathways to the unique conditions in susceptible gut environments following different antibiotic pretreatments (7, 8). These transcriptional shifts indicated that *C. difficile* must coordinate metabolic activity accordingly to compete within new hosts. In spite of these differences, there are known core elements of *C. difficile* metabolism across different environments including carbohydrate and amino acid fermentation (9). It is known that specific growth nutrients influence expression of virulence genes in *C. difficile* (9, 10). Given these findings, targeted therapeutic strategies that alter active metabolism and downregulate virulence may be possible without continued exposure to antibiotics. This form of treatment would be especially beneficial as there have been stark increases in the prevalence of antibiotic resistance and hyper-virulence among *C. difficile* clinical isolates (11, 12).

Genome-scale metabolic network reconstructions (GENREs) are mathematical formalizations of metabolic reactions encoded in the genome of an organism. These models are subsequently constrained by known biological and physical parameters such as membrane transport and enzyme kinetics. GENREs can be utilized to interrogate the metabolic capability of a given organism, as well as providing a means to simulate growth and assess the impact of genotype on metabolism. GENREs have been implemented in directing genetic engineering efforts (13) and accurately predicting auxotrophies and interactions between species for growth substrates (14, 15). These platforms also create improved context for the interpretation of omics data (16), and have provided powerful utility for identification of novel drug and gene targets accelerating downstream laboratory testing (17). This concept is especially critical when delineating a complex array of signals from communities of organisms like the gut microbiome (18). Leveraging these tools, several recent studies have identified nodes of metabolism that promise to provide novel therapeutic targets in clinically-relevant pathogens including *Klebsiella pneumoniae*, *Staphylococcus aureus*, and *Streptococcus mutans* (17, 19, 20). However, there has been limited progress to date applying GENREs to obtain mechanistic understanding for metabolism during infection as they relate to colonization and virulence. Taken together, these principles make GENREs strong platforms for deciphering novel metabolic drivers of virulence-associated phenotypes in *C. difficile*.

We began by generating new GENREs for two strains of *C. difficile* including a highly-characterized laboratory strain *C. difficile* str. 630 (21), as well as a more recently isolated hyper-virulent strain R20291 (22). *De novo* reconstruction for both models was followed by extensive literature-driven manual curation of catabolic pathways and related metabolite transport, with specific emphasis on Stickland fermentation for ATP generation and *C. difficile*-specific redox maintenance (23). Additionally, both GENREs contain a tailored biomass objective function (an *in silico* proxy for bacterial growth, requiring synthesis of major macromolecular components) which accounts for codon biases and amino acid balance, and cell wall structure. Growth simulations from both GENREs were compared against *in vitro* gene essentiality and carbon utilization screens, which indicated significant levels of agreement across all validation datasets.

To assess potential mechanisms of metabolic control of virulence, we then created context-specific models of *C. difficile* metabolism by integrating transcriptomic data collected from both laboratory culture and infection conditions where differential expression of *C. difficile* virulence factors was observed. Overall, during increased virulence expression both strains of *C. difficile* were predicted to favor increased fermentation of amino acids and decreased reliance carbohydrate usage. Specifically in the hyper-virulent strain R20291 during states of phase variation, we found efflux of the biofilm component N-acetylglucosamine in variants known to produce significantly more biofilm experimentally. Additionally, this state was predicted to have increased reliance on glucose to fuel nucleotide synthesis, instead of ATP generation. When tested *in vitro*, we indeed found that the colony morphology associated with this phase variant was dependent on environmental glucose availability. Alternatively in infection-specific models of strain 630, we identified consistent patterns of proline and ornithine fermentation in states of both high and low sporulation, which agreed with metabolomic analysis of each condition. However, in instances of lower spore burden our model predicted significantly greater usage of the host-derived glycan N-Acetylneuraminate and the nucleotide precursor cytidine as primary sources of carbon. In subsequent laboratory testing we were able to show that not only can *C. difficile* use the substrates for growth, but both also lead to lower quantities of spores, which are essential for transmission of the pathogen (24, 25). This work is the first time that contextualized GENREs of a pathogen have been utilized to identify new metabolite signals of virulence regulation. As such, the high-quality GENREs described here can greatly augment the discovery of novel metabolism-directed therapeutics to treat CDI. Moreover, our results demonstrate that GENREs provide an advantage for delineating complex patterns in transcriptomic and metabolomic datasets into tractable experimental targets.

## Results

### *C. difficile* metabolic network generation, gap-filling, and curation

The emergence of hypervirulent strains of *C. difficile* that have unique metabolism and virulence factors highlights the importance for the in-depth study of metabolic pathways to understand and identify targets within these isolates. Core metabolic processes also present an attractive target for novel antimicrobial measures as they may be less likely to allow for acquired antibiotic resistance (26). With these concepts in mind, we focused on the most well-characterized hypervirulent isolate, str. R20291. However, to maximize the utility of the bulk of published *C. difficile* metabolic research, we elected to generate a reconstruction for the lab-adapted str. 630 in parallel. This focus afforded the ability to continuously cross-reference curations between the models and to more readily identify emergent differences that are specifically due to genomic content.

We began the reconstruction process by accessing the re-annotated genome of str. 630 (27) and the published str. R20291 genome (22), both available on the Pathosystems Resource Integration Center database (PATRIC) (28). Following an established protocol for creating high-quality genome-scale models (29), and utilizing the ModelSEED framework and modified reaction database (30), we created scaffold reconstructions for both strains. We subsequently performed complete translated proteome alignment between str. 630 and str. R20291, resulting in 684 homologous metabolic gene products and 22 and 33 unique gene products, respectively (Table S2). Among the distinctive features were additional genes for dipeptides import in str. 630 and glycogen import and utilization in str. R20291, which have both been linked to modulated levels of virulence across strains of *C. difficile* (31, 32).

Manual curation is required to ultimately produce high-quality GENREs and make meaningful biological predictions (33). As such, we proceeded to manually incorporate 259 new reactions (with associated genes and metabolites) and altered the conditions of an additional 312 reactions already present within each GENRE prior to gap-filling (Table S1). Primary targets and considerations for the manual curation of the *C. difficile* GENREs included:

- Anaerobic glycolysis, fragmented TCA-cycle, and known molecular oxygen detoxification (23, 34)
- Minimal media components and known auxotrophies (35-37)
- Aminoglycan and dipeptide catabolism (38-40)
- Numerous Stickland fermentation oxidative and reductive pathways (Table S2) (41-52)
- Carbohydrate fermentation and short-chain fatty acid metabolism (41, 53–55)
- Elements of the Wood-Ljungdahl pathway (56)
- Energy metabolite reversibility (e.g. ATP, GTP, FAD, etc. (57))
- Structural components including teichoic acid, peptidoglycan, and isoprenoid biosynthesis
- Additional pathogenicity-associated metabolites (e.g. p-cresol (44) and ethanolamine (58))

Following the outlined manual additions, we created a customized biomass objective function with certain elements tailored to each strain of *C. difficile*. Our biomass objective function formulation was initially adapted from the well-curated GENRE of the close phylogenetic relative *Clostridium acetobutylicum* (59) with additional considerations for tRNA synthesis and formation of cell wall macromolecules, including teichoic acid and peptidoglycan (Table S1). Coefficients within the formulations of DNA replication, RNA replication, and protein synthesis component reactions were adjusted by genomic nucleotide abundances and codon frequencies to yield strain-specific biomass objective functions (60). To successfully simulate growth, we next performed an ensemble-based pFBA gap-filling approach (61, 62), utilizing a metabolic reaction database centered on Gram-positive anaerobic bacterial metabolism (see Materials & Methods). Gap-filling refers to the automated process of identifying incomplete metabolic pathways due to an absence of genetic evidence that are necessary for *in silico* growth, and addition of the minimal functionality needed to achieve flux through these pathways (63). We performed gap-filling across six distinct and progressively more limited media conditions; complete medium, Brain-Heart Infusion (BHI (64)), *C. difficile* Defined Medium +/- glucose (CDM (37)), No Carbohydrate Minimal Medium (NCMM (5)), and Basal Defined Medium (BDM (35)) (Table S1) which added a total of 68 new reactions that allowed for robust growth across all conditions.

The final steps of the curation process were focused on limiting the directionality of reactions known to be irreversible, extensive balancing of the remaining incorrect reaction stoichiometries, and adding annotation data for all network components. We repeated the assessments that were performed for the earlier reconstructions and found that our GENREs had substantial improvements in all metrics including few, if any, flux or mass inconsistencies and now each received a cumulative MEMOTE score of 86% (Table S1). The new network reconstructions were designated iCdG709 (str. 630) and iCdR703 (str. R20291).

### *C. difficile* GENRE validation against laboratory measurements

A standard measurement of GENRE performance is the comparison of predicted essential genes for growth *in silico* and those found to be essential experimentally through forward genetic screens (65). For a gene to be considered essential, less than 1% of optimal biomass can be produced by a given mutant (the equivalent of no observable growth) during single-gene knockout simulations (66). Recently a large-scale transposon mutagenesis screen was published for str. R20291 (67), and as such we utilized the proteomic alignment from the previous section to determine homologs in str. 630. Simulating growth in BHI rich medium we identified essential genes for both models, which revealed overall accuracies of 89.1% and 88.9%, with negative-predictive values as high as 90.0% for iCdR703 and 89.9% for iCdG709 (Figure S1A). This high degree of agreement supported that metabolic pathways in the new GENREs were structured correctly, and are more likely to provide useful downstream predictions

To then assess if GENRE requirements reflected the components of minimal medium derived experimentally, we identified the minimum subset of metabolites necessary for growth. Through systematic limitation of extracellular metabolites, we were able to determine the impact of each component on achieving some level of biomass flux (Figure S1C). This analysis revealed that most metabolites found to be essential during growth simulation have also been shown experimentally to be required for *in vitro* growth. Interestingly, while growth simulations indicated that neither iCdG709 (str. 630) nor iCdR703 (str. R20291) were auxotrophic for methionine, the published formulation of BDM indicates methionine is found to be largely growth-enhancing but not essential for small levels of growth (36). Additionally, it has been demonstrated in the laboratory that *C. difficile* is able to harvest sufficient bioavailable sulfur from excess cysteine instead of methionine (37, 68), further supporting growth simulation results. Similarly, the published formulation of BDM indicates that pantothenate (vitamin B5) only appears to enhance growth rate *in vitro* and is not necessarily required to support slow growth rates. Our results also indicated that iCdR703 was not auxotrophic for isoleucine relative to iCdG709, and indeed contained additional genes coding for synthesis of a precursor (3S)-3-methyl-2-oxopentanoate (*ilvC*, a ketol-acid reductoisomerase) which were not present in its counterpart GENRE (Table S2). In summary, the *in silico* minimal requirements for iCdG709 and iCdR703 closely mirrored experimental results for both strains of *C. difficile* in the laboratory.

We next assessed additional carbon sources that impact the growth yield predictions for both GENREs. Utilizing previously published results for both *C. difficile* strains in a high-throughput screen (69), we simulated growth for each carbon source individually in background minimal medium and calculated the shift in optimal growth rate. Importantly, *C. difficile* is auxotrophic for specific amino acids (e.g. proline; Fig S1C) that it is also able to catabolize through Stickland fermentation (70), so the background medium must be supplemented with small concentrations of each. As such, the values are reported as the ratio of the final optical density for growth with the given metabolite versus low levels of growth observed in the background medium alone. Through correlation of the results from these two comparisons, we were able to assess how well *in silico* predictions matched experimental results. Across all the 116 total metabolites that were in both the *in vitro* screen as well as the *C. difficile* GENREs, we identified significant predictive correlations in the amount of growth enhancement for iCdG709 and iCdR703 (*p*-values < 0.001) (Figure 1A & 1B). This relationship was even more pronounced for carbohydrates and amino acids, the primary carbon sources for *C. difficile*. When these predictions were reduced to binary interpretations of either enhancement or non-enhancement of growth, we found that iCdG709 predicted 92.8% and iCdR703 predicted 96.6% true-positive enhancement calls (Figure S1B). This metric is most valuable here as it indicates that each GENRE possesses the necessary machinery for catabolizing a given metabolite. Collectively, these data strongly indicated that both GENREs were well-suited for prediction of growth substrate utilization in either strain of *C. difficile*.

**Figure 1).**
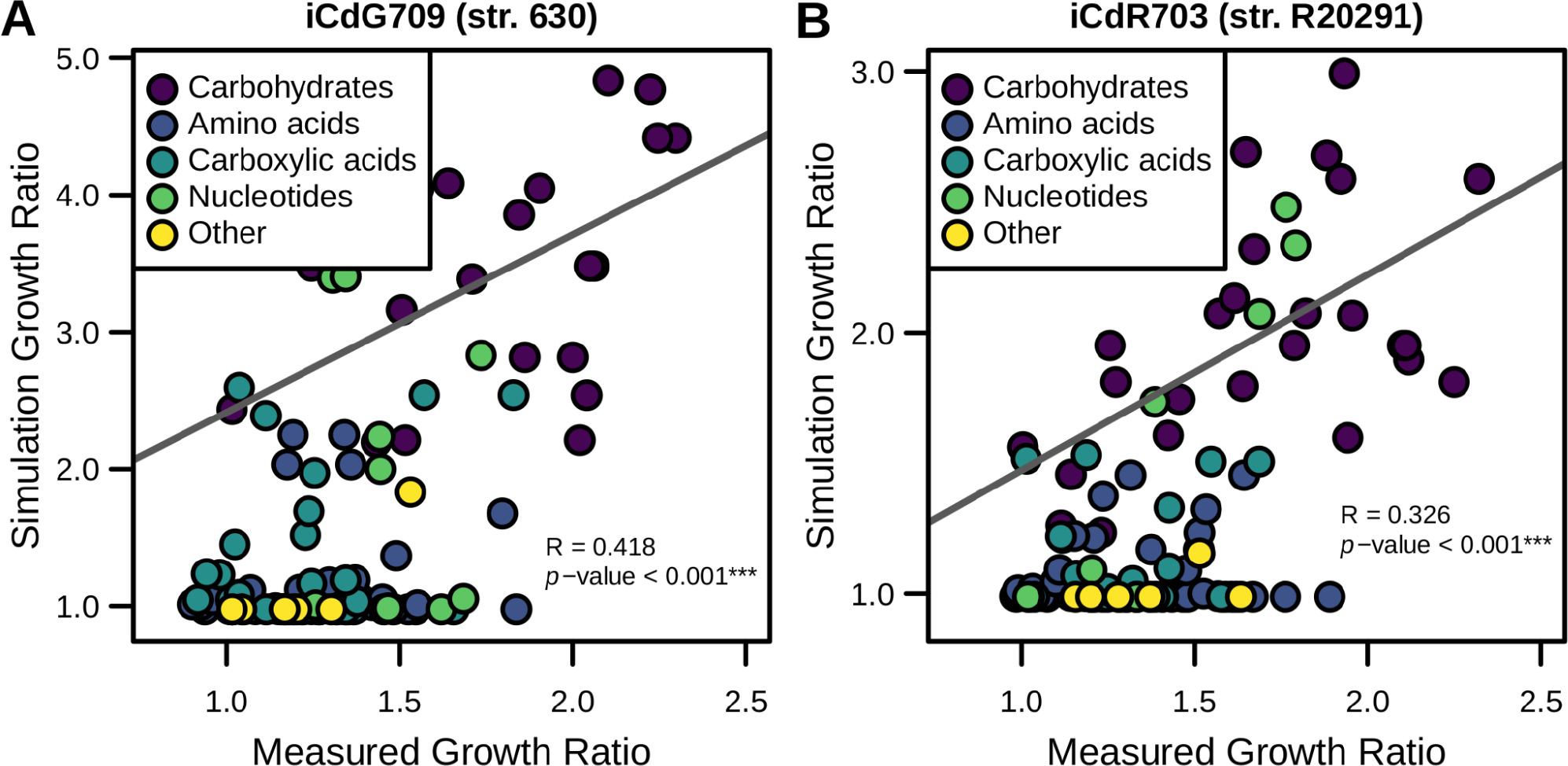
Carbon source utilization prediction profiles accurately reflect laboratory measurements. Results from previous phenotypic screen of 115 metabolites for both str. 630 and str. R20291 were compared against *in silico* results for each corresponding GENRE. Ratios of overall *in vitro* growth enhancement by each metabolite were correlated with the corresponding results from growth simulations in the same media for **(A)** iCdG709 (str. 630) and **(B)** iCdR703 (str. R20291). Points are colored by their biochemical grouping, fit and significance determined by Spearman correlation.

Finally, we also compared our results against existing *C. difficile* GENREs. The primary focus of curated *C. difficile* metabolic modeling efforts has been on the first fully sequenced strain of *C. difficile*, str. 630. The first reconstruction effort (iMLTC806cdf (71)) and subsequent revision (icdf834 (71, 72)), were followed by a recent *de novo* creation following updated genome curation (iCN900 (73)) (27). Another GENRE was developed for str. 630Δerm (iHD992 (74)), a strain derived from str. 630 by serial passage until erythromycin resistance was lost (75). Four additional *C. difficile* strain GENREs were generated as a part of an effort to generate numerous new reconstructions for members of the gut microbiota (76); these reconstructions received only semi-automated curation performed without *C. difficile*-specific considerations. To establish a baseline for the metabolic predictions possible with current *C. difficile* GENREs, we selected common criteria with large impacts on the quality of subsequent predictions for model performance (Table S3). The first of these metrics is the level of consistency in the stoichiometric matrix (57, 77, 78), which reflects proper conservation of mass and that no metabolites are incorrectly created or destroyed during simulations. The next metric is a ratio for the quantity of metabolic reactions lacking gene-reaction rules to those possessing associated genes (79), which may indicate an overall confidence in the annotation of the reactions. These features reflect the importance of mass conservation and deliberate gene/reaction annotation which each drive confidence in downstream metabolic predictions, omics data integration, and likelihood for successful downstream experimentation. We found unique challenges within each GENRE which made comparing simulation results across models difficult. Neither iMLTC806cdf nor iHD994 have any detectable gene annotations associated with the reactions they contain. A high degree of stoichiometric matrix inconsistency was detected across icdf834, iHD992, and iCN900; with iHD992 many intracellular metabolites were able to be generated without acquiring necessary precursors from the environment. We also detected structural inconsistencies across several GENREs. For example, those GENREs acquired from the AGORA database possessed several intracellular metabolic products not adequately accounted for biologically (Table S3), as well as mitochondrial compartments despite being bacteria. Additionally, several key *C. difficile* metabolic pathways either were incomplete or absent from the curated models including multi-step Stickland fermentation, membrane-dependent ATP synthase, dipeptide and aminoglycan utilization, and a variety of saccharide fermentation pathways (23). Considering each of these factors, the *C. difficile* GENREs generated here correct numerous mass and annotation inconsistencies, contain key functional capacities, and phenotypically mimic *C. difficile*.

### Context-specific modeling to capture virulence-associated metabolism

Following validation, we sought to utilize each GENRE to predict *in situ* metabolic phenotypes that correspond with expression of known virulence traits in *C. difficile*. As previously stated, GENREs have provided powerful platforms for the integration of transcriptomic data, creating greater context for the shifts observed between conditions and capturing the potential influence of pathways not obviously connected (80). With this application in mind, we chose to generate context-specific models for both *in vitro* and *in vivo* experimental conditions characterized with RNA-Seq analysis utilizing a recently published unsupervised transcriptomic data integration method (18). Briefly, the algorithm calculates the most cost-efficient usage of the metabolic network to achieve growth given the pathway investments indicated by the transcriptomic data. This approach is in line with the concept that natural selection generally selects against wasteful production of cellular machinery (81). The output models contain only those metabolic reactions that are most likely to be active under the given conditions, whose ranges of metabolic reaction activity were subsequently deeply sampled to assess for distinct yet equally optimal combinations of active pathways. Analysis of these distributions affords the ability to make much more fine-scale predictions of metabolic changes that *C. difficile* undergoes as it activates pathogenicity. The patterns of active pathways also reveal critical elements within context-specific metabolism that could lead to targeted strategies for intentional downregulation of virulence factors through metabolite-focused interventions.

### Phase variation in *C. difficile* str. R20291 is sensitive to carbohydrate availability

*C. difficile* is known to utilize phase variation, a reversible mechanism employed by many bacterial pathogens to generate phenotypic and metabolic heterogeneity to maximize overall fitness of the population. Phase variation has been shown to also influence virulence expression in *C. difficile* str. R20291 (82). One aspect of this phase variation manifests as a rough or smooth-edged colony morphology on solid agar; the morphologies can be propagated via subculture and are associated with distinct motility behaviors and altered virulence (83). The colony morphology variants are generated through the phase variable (on/off) expression of the *cmrRST* genes. Toward understanding this phenotype, we experimentally generated rough and smooth phase variants of *C. difficile* str. R20291 grown on solid BHIS rich medium for 48 hours and sequenced transcriptomes from both groups. Utilizing these data, we generated context-specific versions of iCdR703 in simulated rich media conditions and deeply sampled the resultant metabolic flux distributions to assess all possible forms of metabolism given the new constraints.

While it has been previously shown that mutation of *cmr*-family genes does not significantly alter growth rate *in vitro* (83), the contextualized models predicted significantly increased biomass flux generation (reflective of growth rate) with smooth colony-associated metabolism (Figure S2A). This result fits with experimental findings as the rough-edged phenotype only emerges after long periods of incubation on solid agar when growth rate is measurably slowed (43). We moved on to evaluate structural differences between the context-specific models and identified those metabolic reactions predicted to be active in only the Smooth or Rough context-specific model. With this analysis we found 19 reactions that were distinctly active between conditions (Figure 2A). We then calculated median absolute activity for each reaction which indicated the magnitude at which each reaction contributed to optimal growth in each model. This investigation revealed proline or ornithine fermentation were present and active in either model (Figure 2A). *C. difficile* is capable of easily converting ornithine into proline (52), which is subsequently fermented to 5-aminovalerate for energy. This finding illustrated that proline Stickland fermentation was an integral part of *C. difficile* metabolism across conditions. The finding that N-acetylglucosamine transport only present within the Smooth variant context-specific model was striking as this phase has been previously associated with significantly increased biofilm formation (83), in which N-acetyl-D-glucosamine is the primary component (84). Observing the predicted reaction activity, N-acetylglucosamine transport was not only present exclusively in the Smooth variant context-specific model, but this reaction was extremely active under these conditions (Figure 2C). Furthermore, efflux of the related metabolite D-glucosamine was also significantly increased in the Smooth model (Figure 2D; *p*-value < 0.001). These results supported that the differences in context-specific model structure seen between phase variants likely represented real variation in active metabolism.

**Figure 2).**
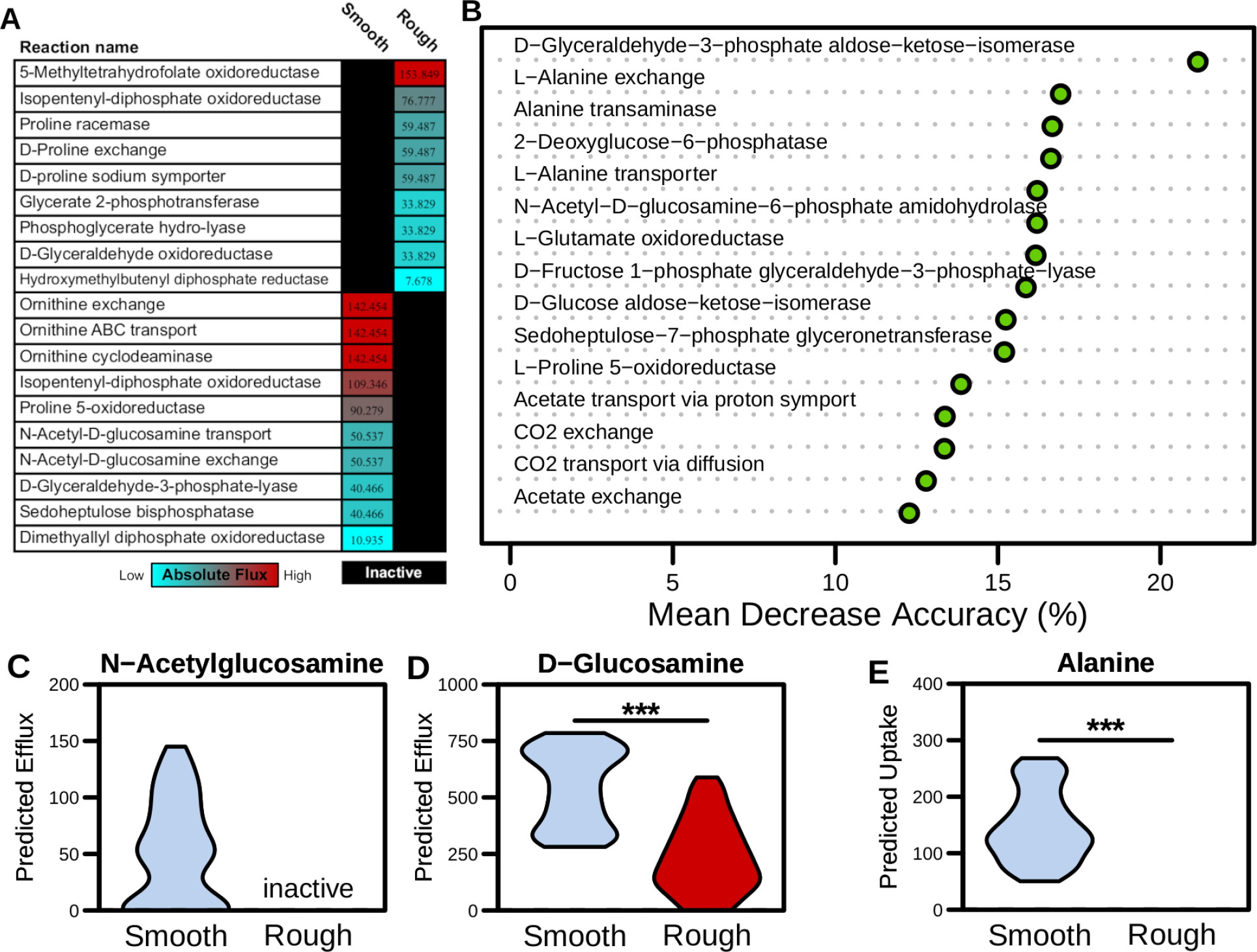
Metabolism significantly varies between phase variants of *C. difficile* str. R20291. Transcriptomes were collected from Rough or Smooth colony morphology clones grown on BHIS agar for 48 hours, and subsequently used to generate context-specific models of *C. difficile* str. R20291. Subnetworks of metabolism that were predicted to be unused in each context were inactivated for subsequent growth simulations. **(A)** Metabolic reactions that are uniquely active in each context-specific model and the associated median absolute reaction activities. **(B)** Utilizing Random Forest supervised machine learning sampled activity for shared non-biomass metabolic reactions between both Rough and Smooth context-specific models (i.e. core metabolism). Shown are the Mean Decrease Accuracy for the top 15 most differentiating reactions. **(C** & **D)** Export exchange reaction flux samples (n = 500) between phase variants for N-Acetylglucosamine and Glucosamine (*p*-value < 0.001). **(E)** Import exchange reaction absolute fluxes between phase variants for Alanine (*p*-value < 0.001). Inactive label denotes reactions pruned during transcriptome contextualization and all significant differences determined by Wilcoxon rank-sum test.

To then compare metabolic activity effectively between context-specific models, we next focused our analysis on shared non-biomass associated reactions across context-specific models which we referred to as “core” metabolism within each subsequent analysis. We first employed unsupervised machine learning for flux samples from core reactions using Non-Metric Multidimensional Scaling (NMDS) ordination of Bray-Curtis dissimilarities (Figure S2B). This analysis revealed significantly different patterns of core metabolic activity between smooth and rough context-specific models (*p*-value = 0.001). To further explore the specific differences within active metabolism between phase variants, we utilized a supervised machine learning approach with Random Forest to discriminate between Rough and Smooth core metabolic activity (Figure 2B). Several of the metabolic reactions with highest mean decrease accuracies are involved in alanine transport and utilization. Further examination of alanine transport reaction fluxes revealed that import and utilization of alanine was only predicted in the Smooth context (Figure 2E). Alanine has been previously identified as having a strong impact on *C. difficile* life cycle physiology (85), and has also been shown to be essential for proper biofilm formation in other Gram-positive pathogens (86). Our results indicate that utilization of alanine may also play a role in biofilm formation and phase variation in *C. difficile*.

Both the network topology and metabolic activity-based analyses indicated that a large number of reactions relating to glycolysis were differentially active. To more closely investigate the relative importance of these metabolic pathways between phase variants, we performed gene essentiality analysis for both models and cross-referenced the results for metabolic reactions associated with the uptake and utilization of glucose (Fig 3A). Through this comparison, we found numerous reactions that were essential only in the Smooth context-specific model which included multiple steps in the Pentose Phosphate Pathway (involved in nucleotide synthesis and NADPH balance) as well the reactions bridging Glycolysis with Fatty Acid Synthesis. Strikingly, no reactions in either pathway were found to be uniquely essential in the rough context-specific model. Although some components of Glycolysis were essential in both contexts, including pyruvate kinase, the penultimate step with the bulk of the ATP production, was detected at the transcriptional level at nearly identical levels between the rough and smooth isolates (Table S4). These findings together signified that ATP generation from Glycolysis was important in both contexts, but the nucleotide precursors and redox potential generated from the Pentose Phosphate Pathway were necessary for the Smooth variant-specific metabolism. In line with this observation, the Rough context-specific model indeed generated a greater fraction of NADH from Stickland fermentation (Table S4). Based on these data, we hypothesized that this additional dependence on glucose was critical in the Smooth variants and without glucose colony morphology would transition toward a more Rough phenotype.

To test this hypothesis, we generated colonies of either Rough or Smooth morphology using C. difficile str. R20291, grown anaerobically for 48 hours on BHIS agar (Figure S3A). We found that the hallmark metric of Rough morphology is a significant increase in colony perimeter (Figure 3D), and used this measurement for determining subsequent shifts between the phenotypes. Both phase variants were subcultured onto BDM agar plates both with and without 2 mg/ml glucose (Figure 3B, 3C). Following anaerobic incubation for 48 hours we found that rough variants maintained their morphology across both media, with the rough phenotype even exacerbated on the minimal medium. However, while the Smooth variant largely maintained its colony morphology upon subculture onto BDM + glucose, the colonies became significantly Rough when glucose was absent (Figure 3E). The inverse was also true in that the Rough colonies maintained their morphology in the absence of glucose, but significantly decreased in perimeter on BDM + glucose, appearing more Smooth (Figure 3F). Further subculture of each altered morphology from minimal media back onto rich BHI medium also appeared to support consistent switching between the respective morphologies (Figure S3B). Our data supported that the Smooth phase variants relied on glucose for more than strictly ATP generation, and that the Rough morphology is apparent only after extended incubation when *C. difficile* may be locally activating starvation responses and switching toward alternative energy sources. Additionally, when glucose is available *C. difficile* will opt to generate redox potential more efficiently through the Pentose Phosphate Pathway. Furthermore, these results are consistent with the hypothesis that carbohydrate availability impacts phase variation in *C. difficile,* and that environmental stress due to limited nutrients may be a key factor in driving the shift between phases.

**Figure 3).**
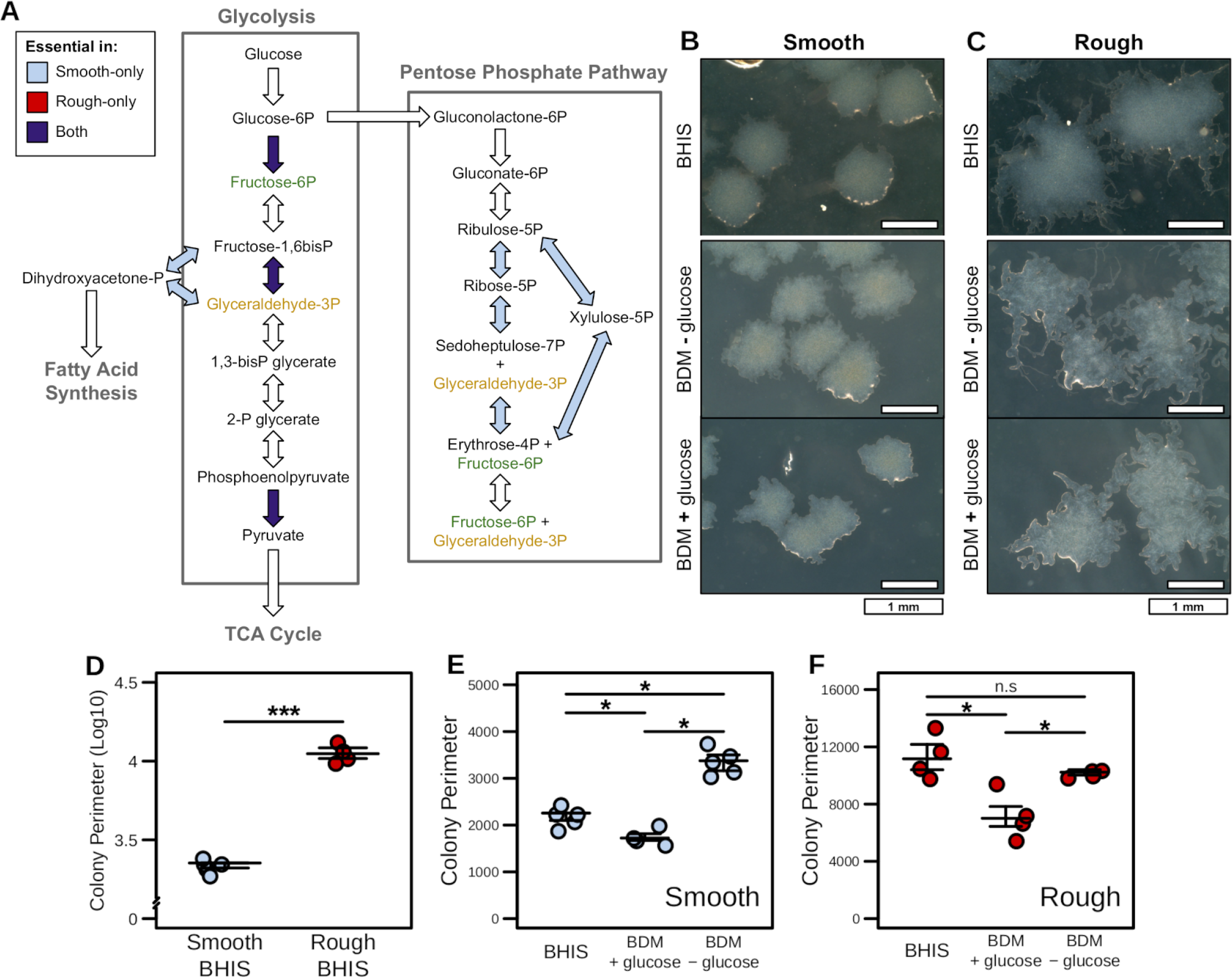
Glucose utilization through the Pentose Phosphate Pathway is essential in the Smooth phase variants of str. R20291. **(A)** Gene and reaction essentiality results for Glycolysis and the Pentose Phosphate Pathway across both the Rough and Smooth phase variant context-specific models. Components were deemed essential if models failed to generate < 1% of optimal biomass flux. **(B & C)** Colony morphologies resulting from smooth and rough variants of *C. difficile* str. R20291 grown on either BHI or BDM +/- glucose (2 mg/ml) after 48 hours of growth (Phase contrast 20/40, 4X magnification). Defined medium colonies were then subcultured onto BHI medium for an additional 24 hours as indicated. Increased colony perimeter was found to be the defining characteristic of the Rough colony morphology. This feature was quantified for multiple colonies under each permutation of colony variant and growth medium (n ≥ 4). **(D)** Colony perimeter for Smooth and Rough progenitor colony variants grown on BHIS (*p*-value < 0.001). **(E)** Smooth or **(F)** Rough colony variant perimeter during subculture onto each of the BDM carbon source media formulations (*p*-values < 0.05). Significant differences determined by Wilcoxon rank-sum test with Benjamini-Hochberg correction when necessary.

### Utilization of N-acetylneuraminic acid and cytidine decreases sporulation in *C. difficile* str. 630

While laboratory conditions are highly informative, it is even more critical to examine metabolism for this pathogen during infection as it can more readily lead to novel therapeutic interventions. It has been previously shown that different classes of antibiotics have distinct impacts on the structure of the gut microbiota while inducing similar sensitivity to colonization by *C. difficile* (87). Along these lines, one published study assessed differential transcriptional activity of *C. difficile* str. 630 in the gut during infection in a mouse model pretreated with the antibiotics cefoperazone or clindamycin. Crucially, these treatments resulted in highly dissimilar levels of sporulation (another phenotype linked to *C. difficile* virulence) where cefoperazone had largely undetectable spore Colony forming units (CFUs), clindamycin had significantly higher levels at the same time point (7). These experiments included paired, untargeted metabolomic analysis of intestinal content to correlate the transcriptional activity of metabolic pathways with changes in the abundance of their respective substrates. Included in the analysis were both mock-infected and *C. difficile*-colonized groups (both treated by the respective antibiotics) to extract the specific impact of the infection on the gut environment, making this dataset extremely valuable.

We first compared predicted biomass objective flux in the sampled context-specific flux distributions (Figure S4A) which revealed no significant difference between high and low sporulation conditions. However, ordination analysis, performed as in the previous section, indeed revealed significant differences in predicted core metabolic activity (Figure 4A; *p*-value = 0.001). In agreement with these findings, supervised machine learning analysis indicated numerous differences in reactions associated with metabolizing host-derived glycans and nucleotide precursors (Figure S4B). To focus this assessment on growth substrates that may be differentially impacting the observed levels of sporulation we assessed each context-specific model by sequentially limiting the ability to import or export each extracellular metabolite to 1% of its optimal rate and measured the impact on overall biomass production (Figure 4B & 4C). Paired metabolomic analysis of each metabolite identified this way were then compared within each condition for relative change in concentration following infection, represented as colored squares along the right margin. Many metabolites had no effect on biomass when their exchange rates were limited, and simply rerouted metabolism elsewhere to achieve similar levels of growth which indicated a high degree of metabolic plasticity remaining in each context-specific model. All metabolites highlighted by this analysis that were measured by the metabolomics screen followed the model-predicted directional change in concentration, supporting the hypothesis that *C. difficile* itself is responsible for the observed differences (Figure 4B & 4C). The peptides proline, ornithine, and serine were found to have an impact on the ability to grow across both context-specific models. Of this subset of amino acids, only proline is an auxotrophy and all are usable by *C. difficile* in Stickland fermentation. Following the catabolism of proline, its Stickland fermentation byproduct 5-aminovalerate was predicted to be an important efflux metabolite in both conditions and had concordant significant increases in concentration following infection in each group (Figure S4C). Alternatively, isovalerate efflux was found to only be critical in higher sporulation context (Figure 4B). This short-chain fatty acid has been primarily associated with leucine fermentation in *C. difficile*, supporting an elevated dependence on Stickland fermentation as sporulation increases. Intestinal concentrations of leucine have indeed been shown to significantly decrease following infection by *C. difficile in vivo* (7), supporting its importance during infection. The most distinguishing features were the importance of N-acetylneuraminate (Neu5Ac) and cytidine only in the lower sporulation context-specific model (Figure 4C). N-acetylneuraminate is a host-derived component of sialic acid that *C. difficile* readily uses as a carbon source for growth (7), and cytidine is an integral component of RNA synthesis. However, neither had been previously associated with directly influencing virulence factor expression in *C. difficile*. Furthermore, while N-acetylneuraminate significantly decreases during infection in the lower sporulation context (7), cytidine also appears to decrease under these conditions implying consumption by *C. difficile* (Figure S4D & S4E).

**Figure 4).**
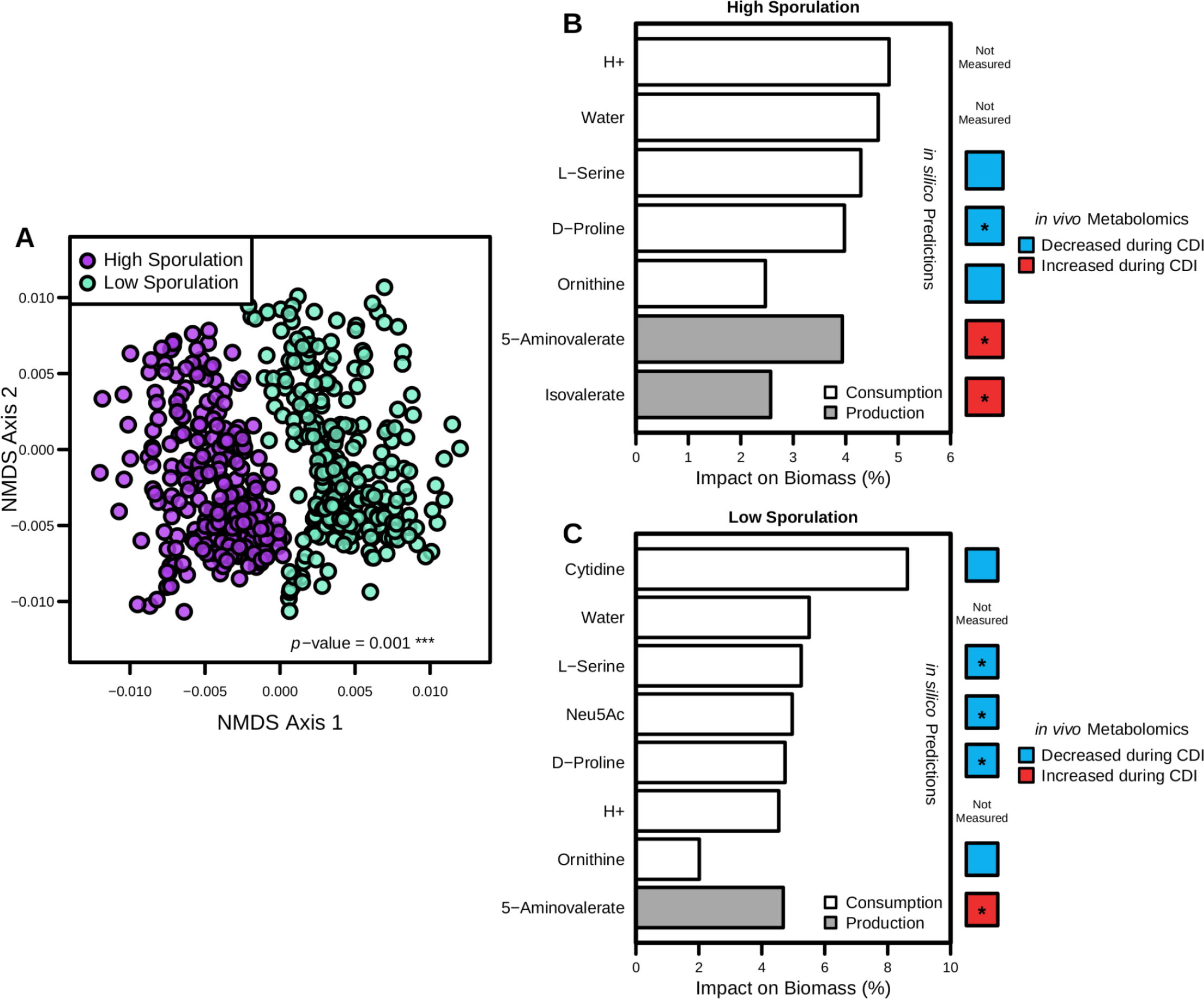
Predicted differences in *C. difficile* str. 630 carbon source usage correspond with lowered rates of sporulation. Transcriptomic integration and predictions with iCdG709, 18 hours post-infection with str. 630 across infections with either high or low levels of sporulation were detected in the cecum. **(A)** NMDS ordination of Bray-Curtis dissimilarities for flux distributions shared reactions following sampling of context-specific models. Significant difference calculated by PERMANOVA. Iterative growth simulations for **(B)** higher sporulation context-specific model and **(C)** in the lower sporulation context-specific model, displaying metabolites with any impact on biomass production when consumption or production capability was restricted to 1.0% of optimal in a given context-specific model. Along the right margin is paired-LCMS analysis from cecal content of mice with and without *C. difficile* str. 630 infection in antibiotic pretreatment groups that resulted in either high or low cecal spore CFUs for metabolites highlighted by growth simulation analysis. Each is colored by mean decrease/increase in concentration between mock and infected groups, and stars indicate significant differences determined by Wilcoxon rank-sum test (*p*-values ≤ 0.05).

We first sought to measure if *C. difficile* str. 630 could utilize both N-acetylneuraminate and cytidine as carbon sources, and if together they exerted a combined effect on growth. Both N-acetylneuraminate and cytidine were supplemented (10 mg/ml each) in liquid BDM in parallel with liquid BDM with no additional substrate and BDM + D-glucose (10 mg/ml) controls, into which *C. difficile* str. 630 was inoculated and incubated for 18 hours and OD600 was measured every 5 minutes (Figure 5A). This assay revealed that *C. difficile* str. 630 could indeed use N-acetylneuraminate and cytidine as carbon sources independently, as each condition allowed for significantly more growth than background BDM alone (*p*-values < 0.05). Additionally, there was no discernible effect on growth when both substrates were added simultaneously. Utilization of the nucleotide precursor cytidine as a carbon source during infection has never been previously described in *C. difficile*, which further supported the utility of our models as a platform for augmenting discovery.

**Figure 5).**
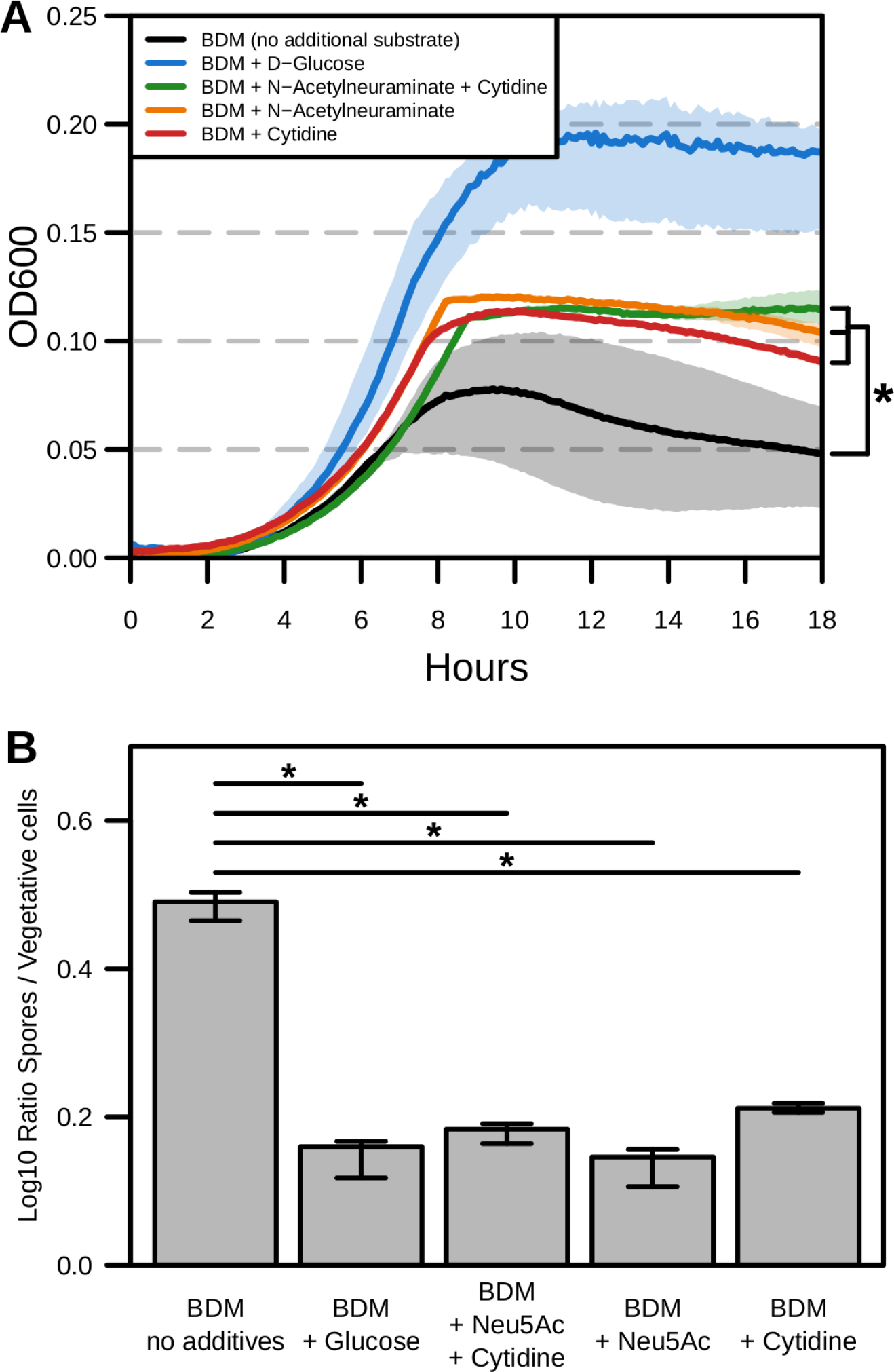
N-Acetylneuraminic acid and cytidine drive changes in str. 630 growth and sporulation. **(A)** 18-hour anaerobic *C. difficile* str. 630 growth measured at OD600 in defined minimal media (BDM) formulated with the indicated carbon sources (10 mg/mL each; n = 4). Significant differences determined using PERMANOVA of Dynamic Time Warping distances (*p*-values < 0.05). **(B)** Median and IQR for the log-transformed *C. difficile* str. 630 spore to vegetative cell CFU ratio after 18-hour incubation in rich medium or defined minimal media (BDM) formulated with the indicated carbon sources (n = 4). Differential plating performed on BHIS-agar +/- taurocholate (1.0%). Significance determined by Wilcoxon rank-sum test with Benjamini-Hochberg correction (*p*-values < 0.05).

To then assess the effect of N-acetylneuraminate and cytidine on sporulation in *C. difficile* str. 630, we performed a growth and sporulation assay targeting these substrates using the same defined medium conditions described previously (Figure 5B). Following 18 hours of anaerobic growth, cultures were plated on BHIS-agar plates lacking a germination agent to quantify specifically vegetative cell CFU abundance. Remaining liquid cultures were then treated with a final concentration of 50% ethanol for 60 minutes to eliminate vegetative cells, and then plated on BHIS-agar with 1% taurocholate added to quantify exclusively spore CFU. The resultant abundances were then converted to an overall spore to vegetative cell ratio to suggest the fraction of the population undergoing sporulation. After overnight incubation, the group that received any combination of N-acetylneuraminate or cytidine had significantly decreased levels of sporulation ratios relative to the no additive control (*p*-values < 0.05), but no significant change when compared to the glucose-added control (Figure 5B). Importantly, there were significantly more vegetative cells in all additive conditions relative to BDM alone (Figure S5; *p*-values < 0.05). As was the case in the growth curve results, there was no difference between N-acetylneuraminate and cytidine when added alone versus their combined effect. Collectively, these results support that both N-acetylneuraminate and cytidine utilization by *C. difficile* inhibit progression through its lifecycle toward spore formation. More broadly, our results support these GENREs as an advantageous discovery platform for novel elements of *C. difficile* metabolism and physiology.

## Discussion

The control for much of *C. difficile*’s physiology and pathogenicity is subject to a coalescence of metabolic signals from both inside and outside of the cell. Historically, *C. difficile* research has suffered from a shortage of molecular tools and high-quality predictive models for highlighting new potential therapies. Over the previous decade, GENREs have become powerful tools for connecting genotype with phenotype and provided platforms for defining novel metabolic targets in biotechnology and improving interpretability of high-dimensional omics data. These factors make GENRE-based analyses extremely promising for directing and accelerating identification of possible therapeutic targets as well as a deeper understanding of the connections between *C. difficile* virulence and metabolism. Furthermore, as much of bacterial pathogenicity is now being attributed to shifts in metabolism the analyses described here may provide large benefits to the identification of possible treatment targets in *C. difficile* and other recalcitrant pathogens (88). In the current study, we develop and validate two highly-curated genome-scale metabolic network reconstructions for a well-described laboratory strain (str. 630) in addition to a more recently characterized hyper-virulent strain (str. R20291) of *C. difficile*. Validation results from both models indicated significant agreement with both gene essentiality and carbon utilization screens, indicating a high degree of confidence in subsequent predictions for active metabolism.

We next employed a recently published technique for transcriptome contextualization with datasets from *in vitro* and *in vivo* systems to evaluate potential emergent metabolic drivers of virulence. These combined analyses revealed differential reliance on glycolysis-related metabolism during periods of increased virulence expression. Specifically, in states of elevated biofilm formation *C. difficile* str. R20291 we found that glucose is necessary for nucleotide synthesis and redox balance through the Pentose Phosphate Pathway, despite still being utilized for ATP in conditions associated with reduced biofilm. These findings were subsequently supported by direct testing *in vitro*, and agreed with recent work which supports that access to glycolysis intermediates actually induces *C. difficile* biofilm formation (89). Alternatively, during infection with str. 630 we identified patterns of host-derived glycan (N-acetylneuraminate) utilization in combination with consumption of the nucleotide precursor cytidine that corresponded with lower levels of sporulation. While not typically considered a carbon source for *C. difficile*, laboratory testing confirmed that *C. difficile* can indeed use cytidine for energy, and along with N-acetylneuraminate, decreases in sporulation. Intentional control of sporulation is an exciting prospect as spores are considered the transmissive form of *C. difficile*, so these results may prove valuable for downstream targeted manipulation of *C. difficile* virulence factor expression. Our results also supported a role for some level of amino acid fermentation across all conditions tested. This phenotype is a hallmark of *C. difficile* physiology and reinforced the validity of the other predictions. These results indicate a complex relationship with environmental nutrient concentrations and likely competition with the gut microbiota that all inform the regulation of *C. difficile* virulence expression. Additionally, *in vivo* context-specific gene essentiality also predicted proline racemase to be critical for growth during infection, yet it was previously found to be dispensable in an animal model using a forward genetic screen (90). This result may be attributable to the specific conditions of that infection and may vary across distinct host gut environments, leading to possible implications in personalized medicine and novel approaches to curbing the expression of virulence factors by influencing environmental conditions to favor certain forms of metabolism over others. This study represents the first time that context-specific models of bacterial metabolism have been generated and used to augment discoveries for metabolic control over virulence expression in the laboratory.

While the majority of predictions followed experimental results, several areas of possible expansion and curation are present in both GENREs. First, while the scope of total genes included in iCdG709 and iCdR703 may be more limited than previous network reconstructions, we elected to focus on those gene sets where the greatest amount of evidence and annotation data could be found to maximize confidence in functionality. Consequently, both GENREs consistently underpredict the impact of some metabolite groups, primarily nucleotides and carboxylic acids, which could be due to the absence of annotation or overall knowledge of the relevant cellular machinery. Furthermore, more complex regulatory networks ultimately determine final expression of virulence factors, and these may be needed additions in the future to truly understand the interplay of metabolism and pathogenicity in *C. difficile*. Despite these potential shortcomings, both iCdG709 and iCdR703 produced highly accurate metabolic predictions for their respective strains as well as novel predictions for metabolism as it relates to *C. difficile* virulence expression, making both strong candidate platforms for directing future studies of *C. difficile* metabolic pathways.

Systems-biology approaches have enabled the assessment of fine-scale changes to metabolism of single species within complex environments that may have downstream implications on health and disease. Overall, the combined *in vitro*- and *in vivo*-based results demonstrated that our GENREs are effective platforms for gleaning additional understanding from omics datasets, outside of the standard analyses. Both GENREs were able to accurately predict complex metabolic phenotypes when provided context-specific omics data, and ultimately underscores the metabolic plasticity of *C. difficile*. The reciprocal utilization of glycolysis and amino acid fermentation indeed support regimes of distinct metabolic programming associated with *C. difficil*e pathogenicity. Finding core metabolic properties in *C. difficile* strains may be key in identifying potential probiotic competitor strains or even molecular inhibitors of metabolic components. The current study is an example of the strength that systems-level analyses have in contributing to more rapid advancements in biological understanding. In the future, the metabolic network reconstructions presented here are well-suited to accelerate research efforts toward the discovery of more targeted therapies. Overall, GENREs have had limited impact to date in real mechanistic understanding of infectious disease and the current study represents a significant advance in this application.

## Materials & Methods

### *C. difficile* GENRE construction

We utilized PATRIC reference genomes from *Clostridioides difficile* str. 630 and *Clostridioides difficile* str. R20291 as initial reconstruction templates for the automated ModelSEED pipeline (28, 91, 92). The ModelSEED draft network reconstruction was converted utilizing the Mackinac pipeline (https://github.com/mmundy42/mackinac) into a form more compatible with the COBRA toolbox (93). Upon removal of GENRE components lacking genetic evidence (i.e. gap-filled), extensive manual curation was performed in accordance with best practices agreed upon by the community (94). We subsequently performed ensemble gap-filling as previously described, utilizing a stoichiometrically consistent anaerobic, Gram-positive universal reaction collection curated for this purpose and available alongside code associated with this study. Next, we corrected reaction inconsistencies and incorrect physiological properties (e.g. ensured free water diffusion across compartments). Final transport reactions were then validated with TransportDB (95). All formulas are mass and charged balanced at an assumed pH of 7.0 using the ModelSEED database in order to maintain a consistent and supported namespace to augment GENRE interpretability and future curation efforts. We then collected annotation data for all model components (genes, reactions, and metabolites) from SEED (94, 96), KEGG (97), PATRIC, RefSeq (98), EMBL (99), and BiGG (100) databases and integrated it into the annotation field dictionary now supported in the most recent SBML version (101). Complete MEMOTE quality reports for both *C. difficile* GENREs are also available in the GitHub repository associated with this study, as well as full pipelines for model generation.

### Growth simulations, flux-based analyses, and GENRE quality assessment

All modeling analyses were carried out using the COBRA toolbox implemented in python (102). The techniques utilized included: flux-balance analysis, flux-variability analysis (103), gapsplit flux-sampler (104), and minimal_medium on exhaustive search settings. GENRE quality assessment tools were also developed in python and are fully available in the project Github repository. MEMOTE quality reports were generated using the web-based implementation found at https://memote.io/.

### *C. difficile* str. R20291 *in vitro* growth and microscopy

*C*. *difficile* str. R20291 growth was maintained in an anaerobic environment of 85% N_2_, 5% CO_2_, and 10% H_2_. The strain was grown on BHIS-agar (37 g/L Bacto brain heart infusion, 5 g/L yeast extract, 1.5% agar) medium at 37 °C for 48 hours to obtain isolated colonies. Rough and smooth colonies were chosen for propagation on BHI-agar to ensure colony morphology maintenance (83). Basal Defined Medium (BDM) was formulated as previously published (35) with the addition of 1.5% agar for plates, and incubated for 48 hours at 37 °C to generate isolated colonies. Microscopy images were taken on an EVOS XL Core Cell Imaging System at 4x magnification. Colony dimensions determined using ImageJ (https://imagej.nih.gov/ij/).

### *C. difficile* str. 630 *in vitro* growth and sporulation assay

*C*. *difficile* str. 630 growth was maintained in an anaerobic environment of 85% N_2_, 5% CO_2_, and 10% H_2_. Liquid BDM formulated as previously described with the indicated combinations of D-glucose (10 mg/ml), N-acetylneuraminic acid (10 mg/ml), and cytidine (10 mg/ml). Overnight BHI liquid cultures of *C*. *difficile* str. 630 were back-diluted 1:3 in fresh anaerobic BHI and incubated for 1 hour at 37 °C at which point 5 µL was inoculated into 1 mL of each media condition. After 18 hours anaerobic incubation at 37 °C, serial dilutions in anaerobic Phosphate Buffered Saline of these cultures were plated on BHIS-agar (37 g/L Bacto brain heart infusion, 5 g/L yeast extract, 1.5% agar) plates to quantify vegetative cell abundance, then treated with 50% EtOH for 30 minutes(105) and serial dilutions in anaerobic Phosphate Buffered Saline were subsequently plated on BHIS-agar + 1.0% taurocholate plates to measure spore abundance. Plates were incubated for an additional 24 hours at 37 °C, at which point CFUs were quantified. For anaerobic growth curves, 250 µL of each medium was inoculated with 5 µL of the back-dilution and the OD600 was measured every 5 minutes for 18 hours (Tecan Infinite M200 Pro).

### RNA isolation, and transcriptome sequencing

For RNA isolation, rough and smooth isolates were subcultured in BHIS broth (37 g/L Bacto brain heart infusion, 5 g/L yeast extract) overnight (16-18 h) at 37 °C, then 5 µL of the cultures were spotted on BHIS agar (1.5% agar). After 24 h, the growth was collected and suspended in 1:1 ethanol:acetone for storage at -20 °C until subsequent RNA isolation. Cells stored in ethanol:acetone were pelleted by centrifugation and washed in TE (10 mM Tris, 1 mM EDTA, pH 7.6) buffer. Cell pellets were suspended in 1 mL Trisure reagent. Silica-glass beads (0.1 mm) were added and cells were disrupted using bead beating (3800 rpm) for 1.5 minutes. Nucleic acids were extracted using chloroform, purified by precipitation in isopropanol followed by washing the cold 70% ethanol, and suspended in nuclease-free water. Samples were submitted to Genewiz, LLC (South Plainfield, NJ, USA) for quality control analysis, DNA removal, library preparation, and sequencing. RNA sample quantification was done using a Qubit 2.0 fluorometer (Life Technologies), and RNA quality was assessed with a 4200 TapeStation (Agilent Technologies). The Ribo Zero rRNA Removal Kit (Illumina) was used to deplete rRNA from the samples. RNA sequencing library preparation was done using the NEBNext Ultra RNA Library Prep Kit for Illumina (NEB) according to the manufacturer’s protocol. Sequencing libraries were checked using the Qubit 2.0 Fluorometer. The libraries were multiplexed for clustering on one lane of the Illumina HiSeq flow cell. The samples were sequenced using a 2x150 paired-end configuration on an Illumina HiSeq 2500 instrument. Image analyses and base calling were done using the HiSeq Control Software. The resulting raw sequence data files (.bcl) were converted to the FASTQ format and de-multiplexed with bcl2fastq 2.17 software (Illumina). One mismatch was permitted for index sequence identification. Data were analyzed using CLC Genomics Workbench v. 20 (Qiagen). Reads were mapped to the *C. difficile* R20291 genome (FN545816.1) using the software’s default scoring penalties for mismatch, deletion, and insertion differences. All samples yielded over 22 million total reads, with over 20 million mapped to the reference (>93% of total reads, and >90% reads in pairs). Transcript reads for each gene were normalized to the total number of reads and gene length (expressed as reads per kilobase of transcript per million mapped reads, RPKM).

### Genomic and transcriptomic data processing

Alignment of *C. difficile* str. 630 and str. R20291 peptide sequences was performed using bidirectional BLASTp. RNA-Seq reads were first quality-trimmed with Sickle with a cutoff ≧Q30 (Joshi & Fass, 2011(106). Mapping curated reads to the respective *C. difficile* genome was then performed with Bowtie2 (107). MarkDuplicates then removed optical & PCR duplicates (broadinstitute.github.io/picard/), and mappings were converted to idxstats format using SAMtools (108). Abundances were then normalized to both read and target lengths. Transcriptomic integration and context-specific model generation were performed with RIPTiDe using the maxfit_contextualize() function on the default settings (18).

### Statistical methods

All statistical analysis was performed in R v3.2.0. Non-metric multidimensional scaling of Bray-Curtis dissimilarity and perMANOVA analyses were accomplished using the vegan R package (109). Significant differences for single reaction flux distributions, metabolite concentrations, spore CFU, and growth over time were determined by Wilcoxon signed-rank test. Supervised machine-learning was accomplished with the implementation of AUC-Random Forest also in R (110). Dissimilarity between C. difficile str. 630 growth curves determined using Dynamic Time Warping (111).

### Data availability

Genomic and proteomic data for the strains *Clostridioides difficile* str. 630 (PATRIC ref. 272563.8) and *Clostridioides difficile* str. R20291 (PATRIC ref. 645463.3) was downloaded from the PATRIC database (91). Transcriptomic data was downloaded in raw FASTQ format from the NCBI Sequence Read Archive (PRJNA415307 and PRJNA354635) and the Gene Expression Omnibus (GSE158225).

### Code availability

Github repository for this study, with all programmatic code and GENREs described here, can be found at: https://github.com/mjenior/Jenior_CdifficileGENRE_2021.

## Author Contributions

MLJ - Conceptualization. Data generation and analysis. Drafting manuscript.

JLL - Conceptualization. Data generation. Editing manuscript.

DAP - Data generation and analysis. Editing manuscript.

EMG - Conceptualization. Data generation. Editing manuscript.

KAW - Data generation. Editing manuscript.

MED - Data analysis. Editing manuscript.

WAP - Supervision. Editing manuscript.

RT - Supervision. Data generation. Editing manuscript.

JP - Funding acquisition. Supervision. Drafting and editing manuscript.

## Acknowledgements

The authors would like to acknowledge Bonnie Dougherty, Laura Dunphy, and Dawson Payne for their input and feedback on modeling parameters and biomass objective function formatting. We would also like to thank Alex Smith and Joe Zackular for discussions on specifics of *C. difficile* metabolism. The authors have declared that no competing interests exist. This work was supported by funding from The U.S. National Institutes of Health awards R01AT010253 to JP, R01Al143638 to RT, to 5R01AI124214 WP, and DK124048-01A1 to JL, as well as a pilot grant from the UVA Trans-University Microbiome Initiative to MJ. The funding agency had no role in study design, data collection/analysis, or preparation of the manuscript.

---

**Figure S1).**
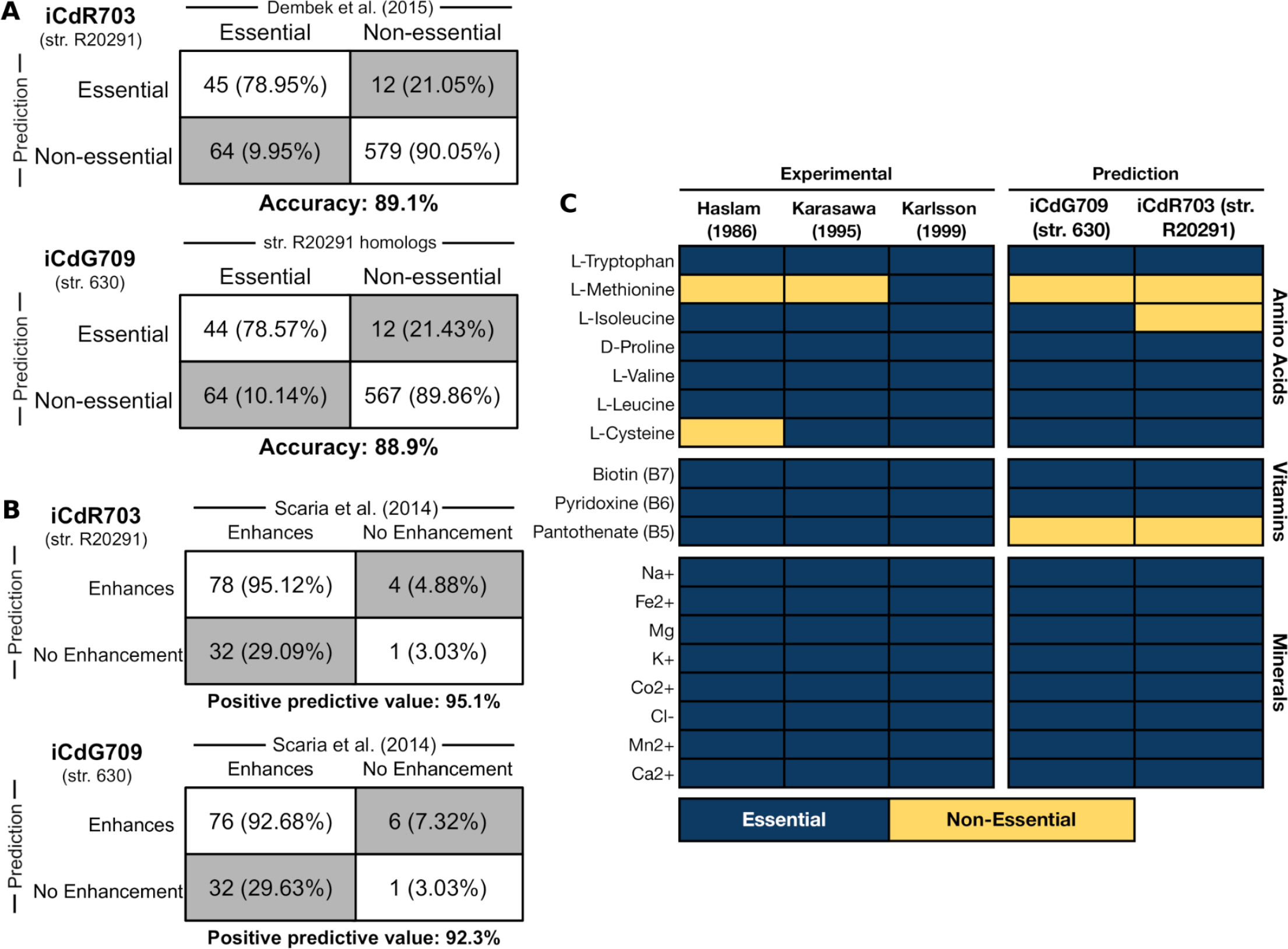
Additional C. difficile GENRE validation against laboratory measurements. **(A)** *in silico* gene essentiality predictions for both GENREs cross-referenced with the Dembek et al. transposon screen for str. R20291 (iCdG709 (str. 630) utilizes homologs from the genome of str. R20291). **(B)** Binary quantification for metabolite growth enhancement shown for both strains/GENREs in Figure 1. Positive predictive values were 95.1% for iCdR703 and 92.3% for iCdG703. **(C)** Computationally determined minimum growth substrates for both GENREs compared with experimentally determined *C. difficile* minimal medium components across three previously published studies. Essentiality was determined for those genes and metabolites that when absent resulted in a yield of <1.0% of optimal biomass flux during growth simulation utilizing components of the corresponding media used experimentally.

**Figure S2).**
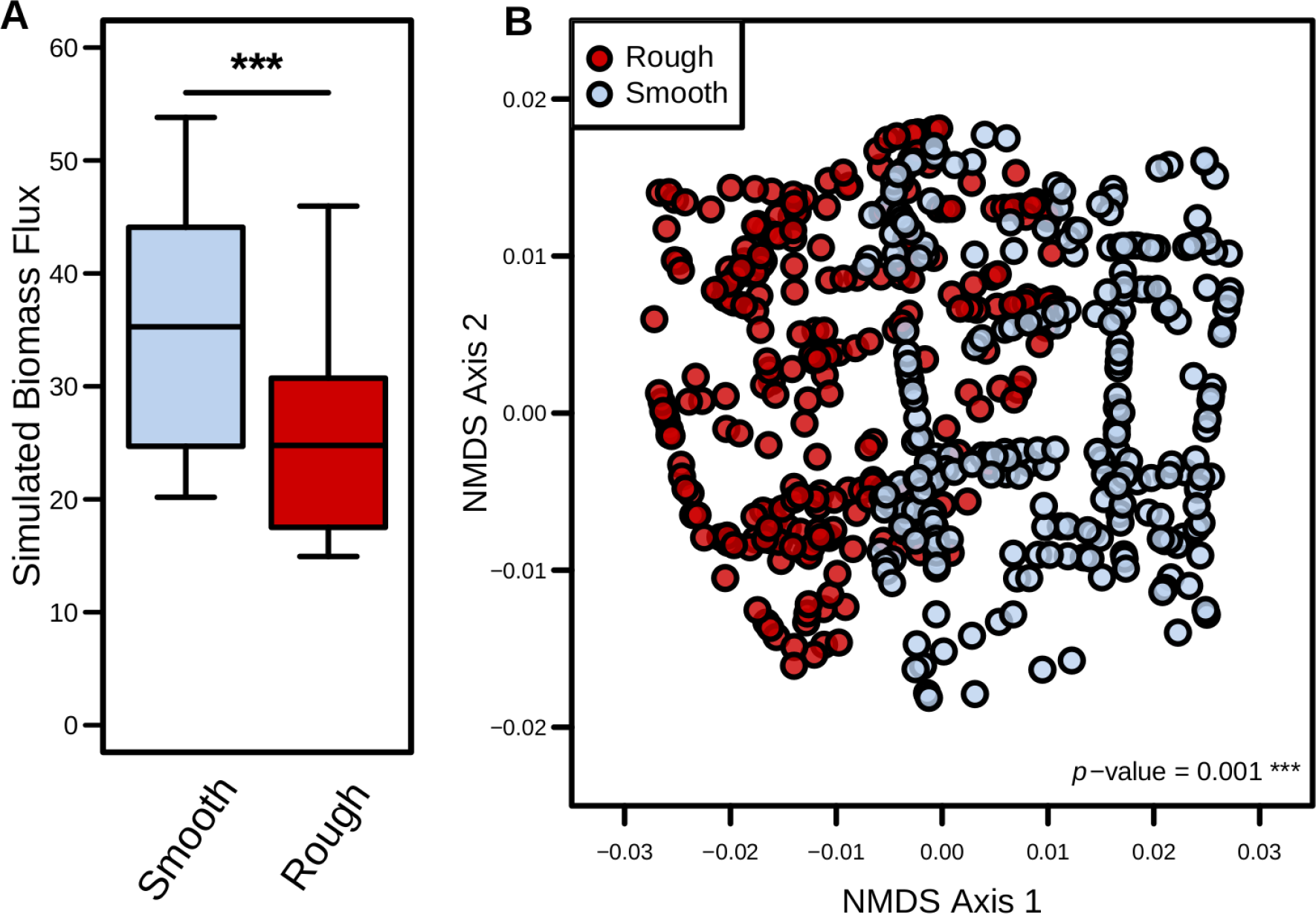
Predicted core metabolic activity significantly differs between str. 630 phase variants. **(A)** Sampled biomass objective flux distributions from each context-specific model. Significance determined by Wilcoxon rank-sum test (*p*-value < 0.001). **(B)** Analysis limited to non-biomass reactions shared across context-specific models of iCdR703. NMDS ordination of Bray-Curtis dissimilarities for flux sampling results from core metabolic reactions. Significant difference determined by PERMANOVA.

**Figure S3).**
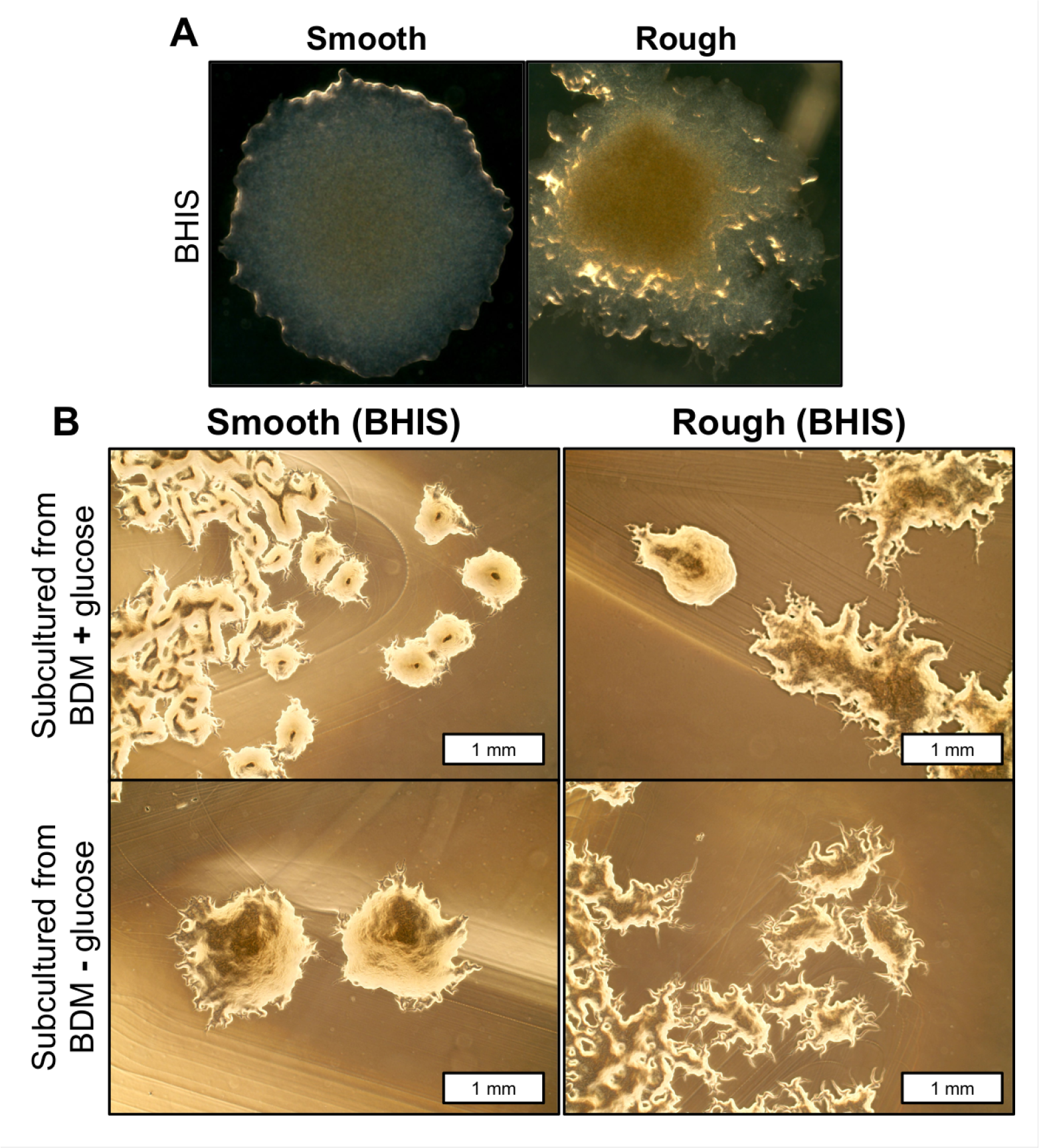
Additional microscopy of phase variant colony morphologies. **(A)** *C. difficile* str. R20291 phase variant progenitor colonies generated on solid BHIS agar following 48 hours of growth at 37° C under anaerobic conditions. These colonies were subcultured and utilized for all subsequent defined minimal medium experiments. **(B)** Subcultured colonies from the indicated conditions in Figure 3B&C onto BHIS rich agar medium, incubated at 37° C for 48 hours anaerobically.

**Figure S4).**
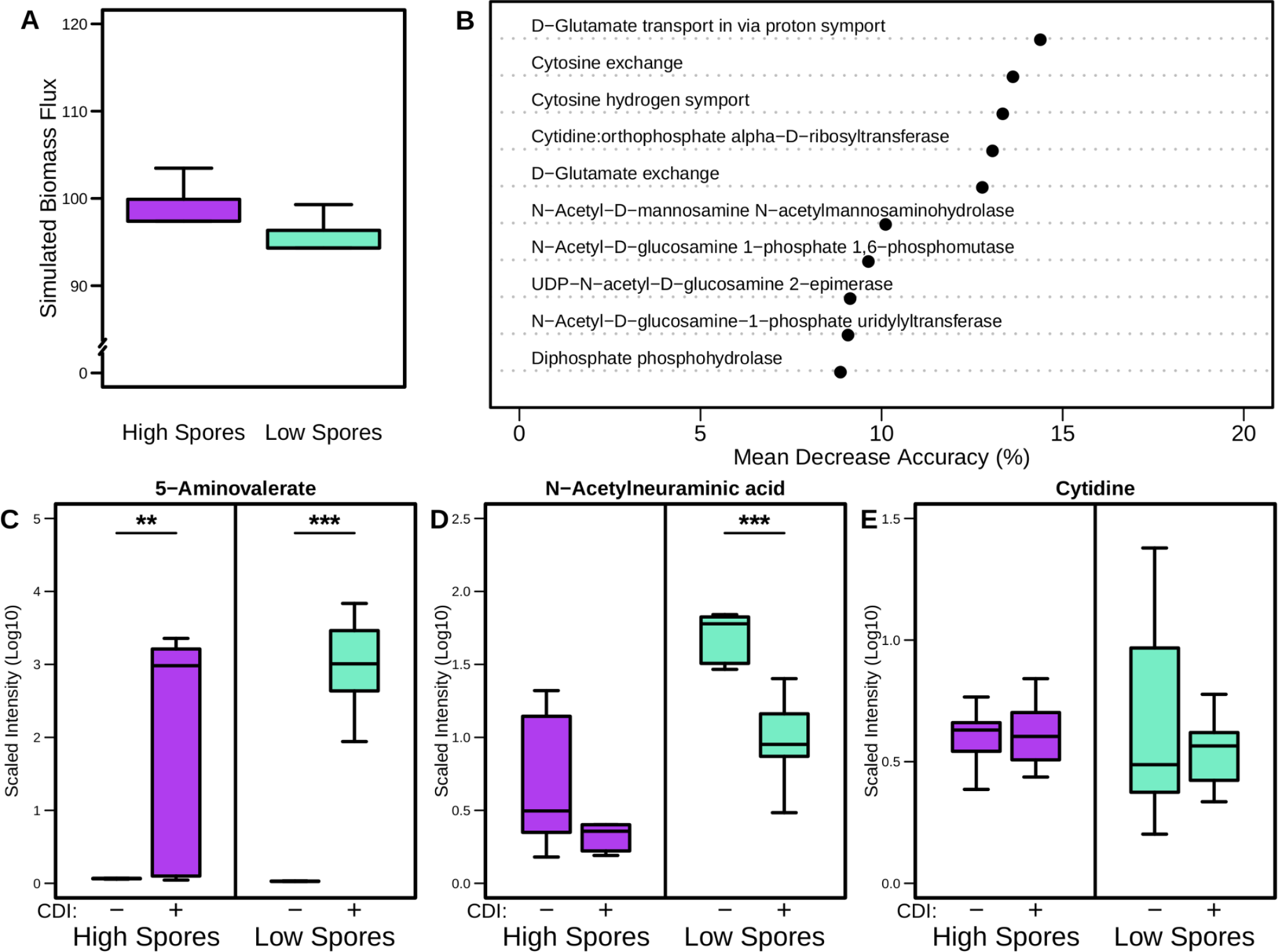
Strain 630 growth simulation and infection metabolomic results. **(A)** Sampled biomass objective flux distributions from each context-specific model. Significant difference tested using Wilcoxon rank-sum test (*p*-value = n.s.). **(B)** Mean Decrease Accuracy for top 10 most differentiating features/reactions from Random Forest supervised machine learning for shared non-biomass reactions across context-specific models of iCdG709. **(C)** Cecal concentrations of 5-aminovalerate in mock and *C. difficile* str. 630 infected mice pretreated with streptomycin, 18 hours after initial colonization. Significance determined by Wilcoxon rank-sum test; *p*-values = 0.003 (High spores) & 0.0001 (Low spores).

**Figure S5).**
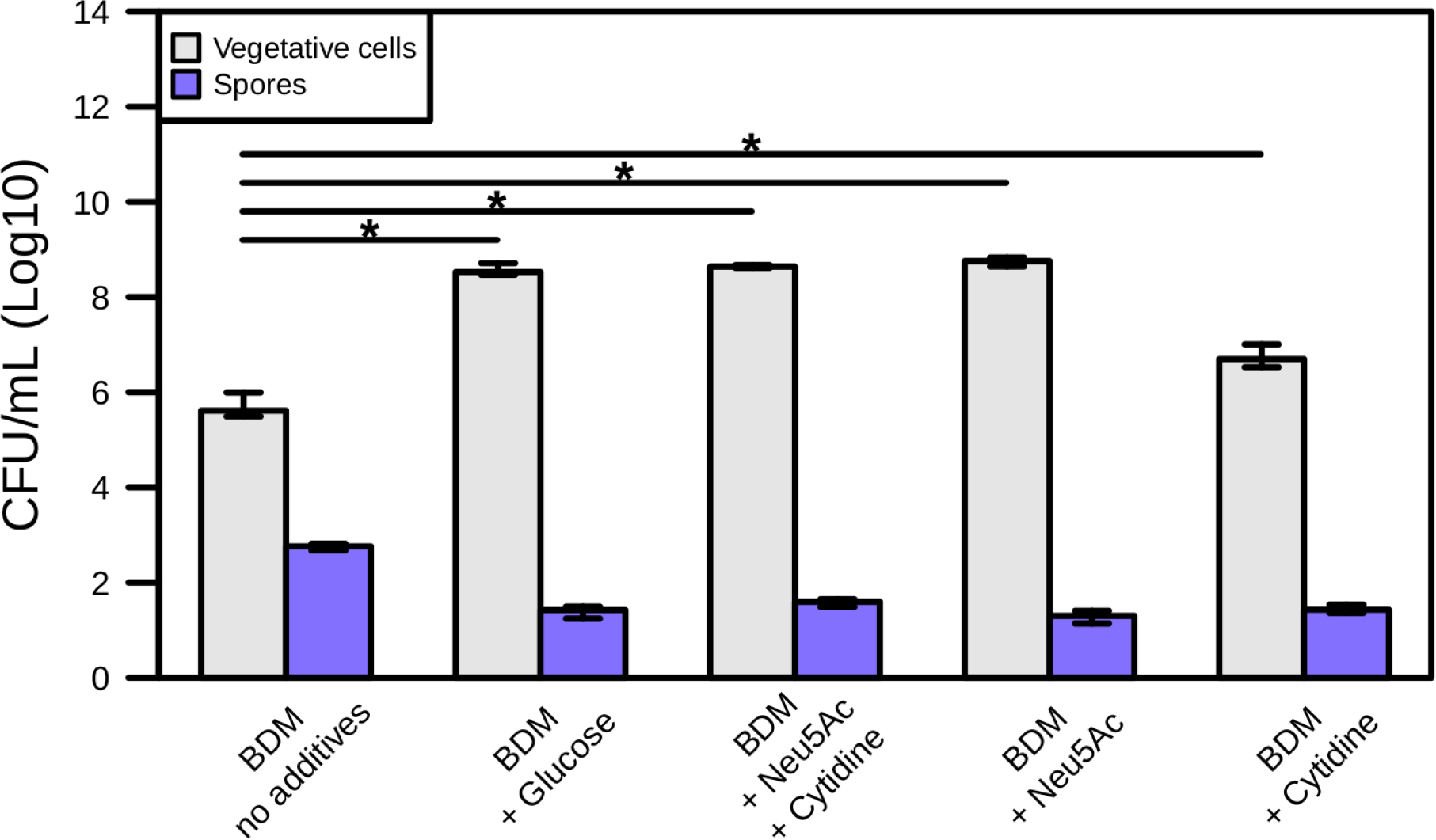
Raw *C. difficile* str. 630 spore and vegetative cell quantification in defined media sporulation assay. Median and IQR of *C. difficile* str. 630 vegetative cells and spore CFU (n = 4) after 18 hour incubation in rich medium or minimal media (BDM) formulated with the indicated additional carbon sources (10 mg/mL each). Significant differences calculated with Wilcoxon rank-sum test with Benjamini-Hochberg correction (*p*-values < 0.05).

**Table S1)** GENRE creation steps, Biomass formulation, Gap-filling media compositions, and GENRE statistics.

**Table S2)** C. difficile 630 and R20291 PATRIC protein sequence alignment results.

**Table S3)** Topology summary statistics for *C. difficile* GENREs from AGORA and those generated here.

**Table S4)** Rough vs Smooth context-specific analysis of iCdR703 (str. R20291)

**Table S5)** High sporulation vs Low sporulation context-specific analysis of iCdG709 (str. 630).

