## Supplementary figures and images for "Novel drivers of virulence in *Clostridioides difficile* identified via context-specific metabolic network analysis"

### Figure S1

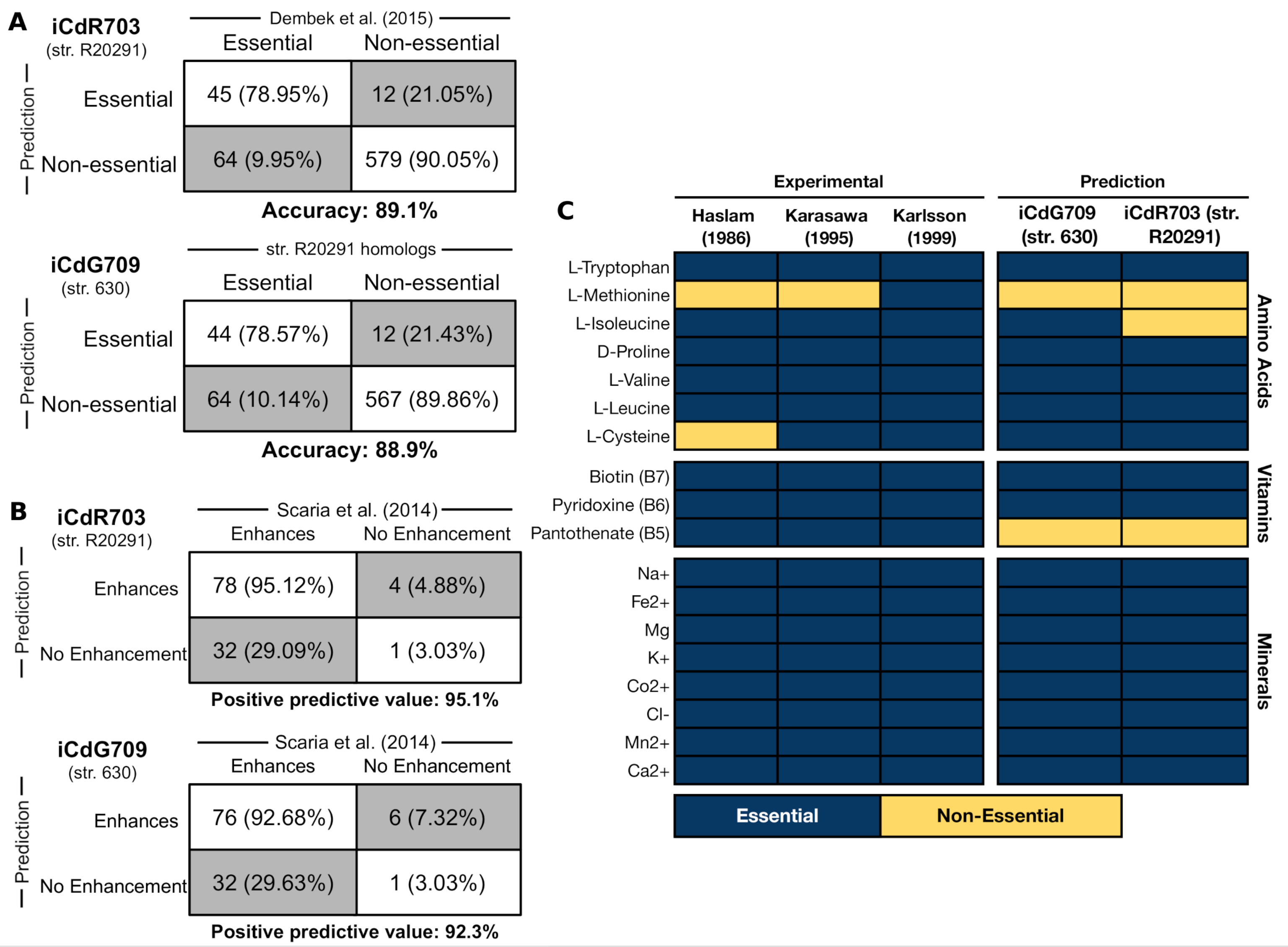

### Figure S2

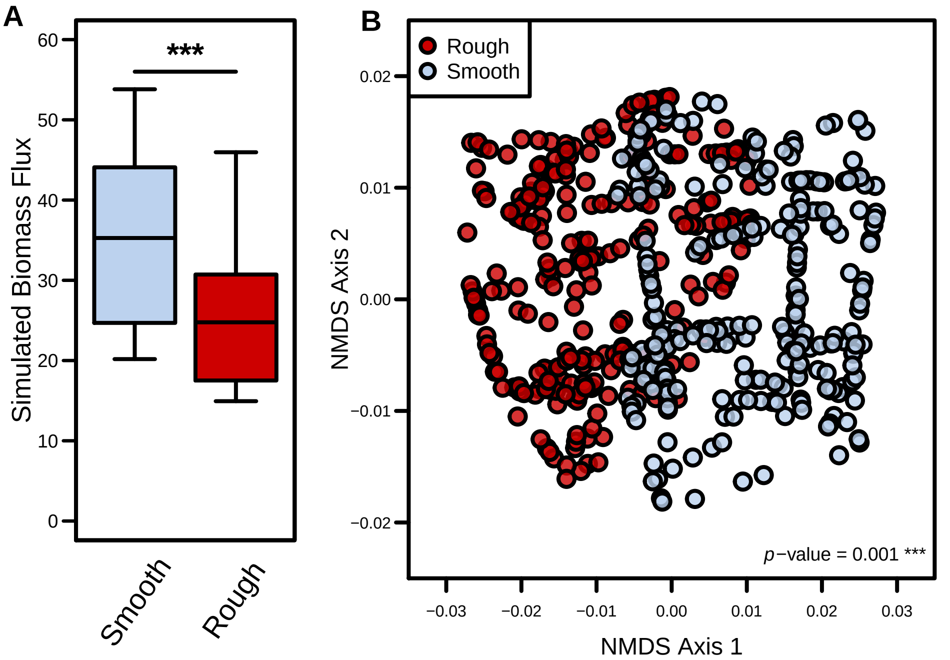

### Figure S3

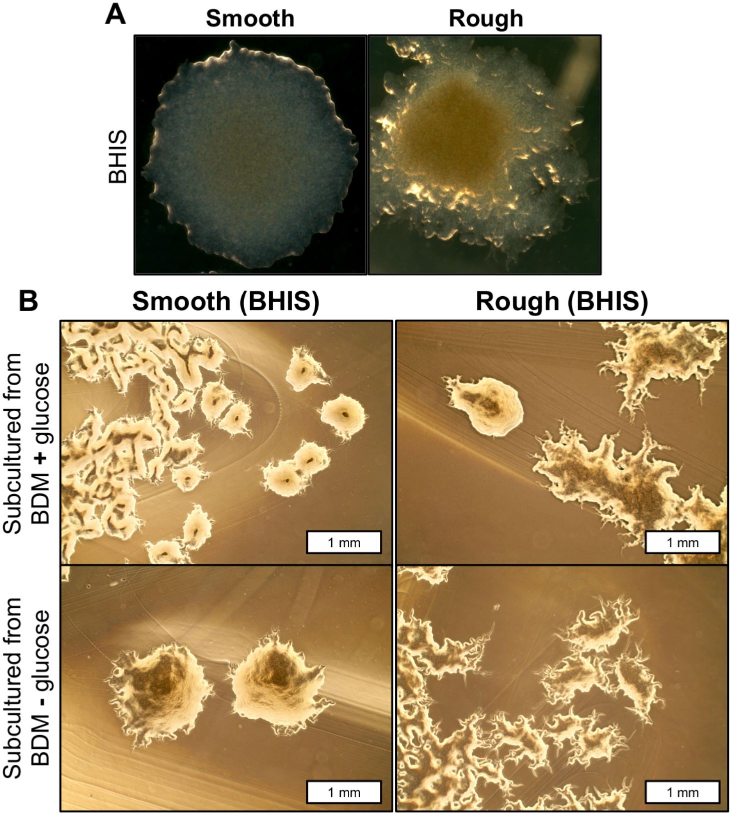

### Figure S4

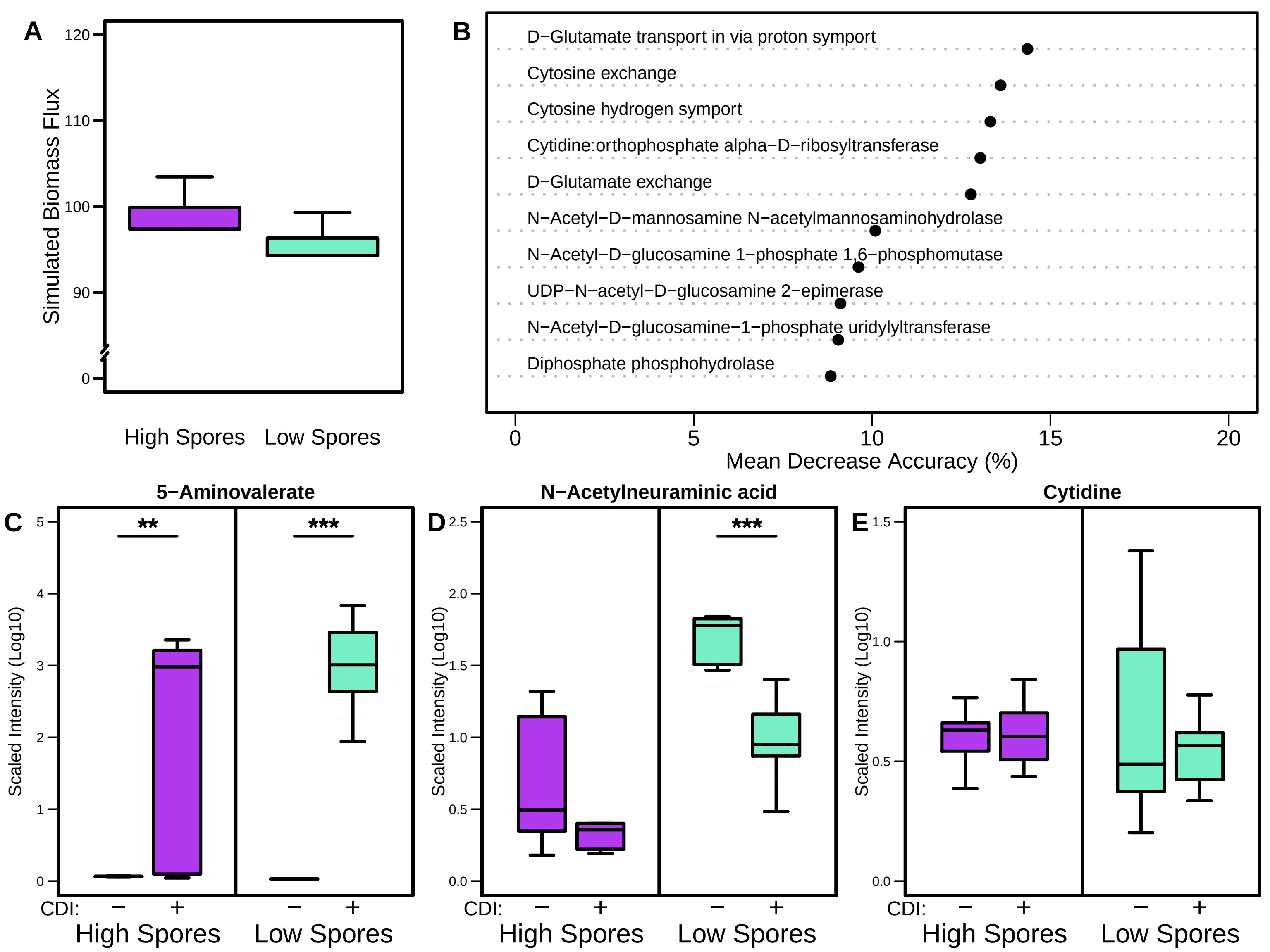

### Figure S5

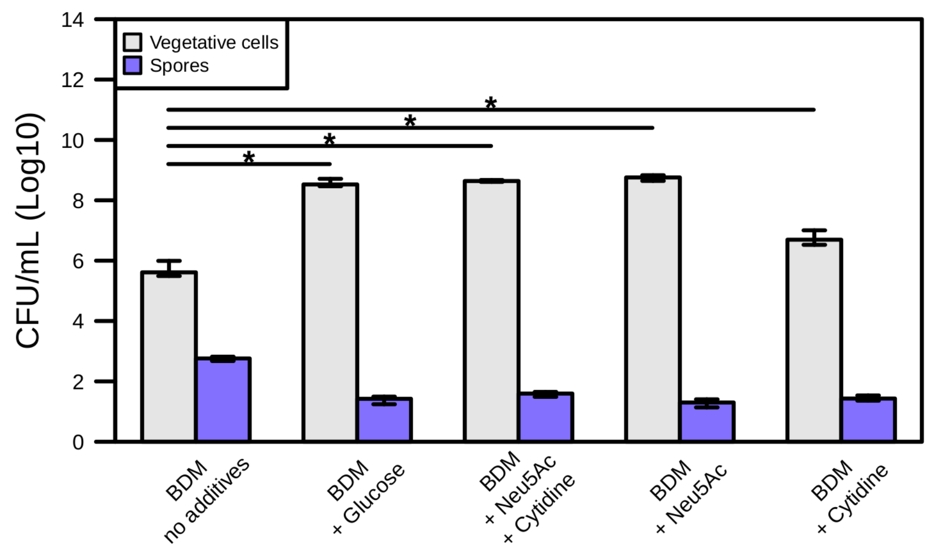
